# Right Caudate Volume and Executive Functions in Children with Attention-Deficit/Hyperactivity Disorder (ADHD)

**DOI:** 10.1101/2024.09.04.611072

**Authors:** Tasmia Hai, Rose Swansburg, Cynthia K. Kahl, Martin Mrazik, Jean-François Lemay, Frank P. MacMaster

**Affiliations:** Department of Educational and Counselling Psychology, McGill University, 3700 McTavish Road, Education Building, Montreal, QC, H3A 1Y9, Canada; Werklund School of Education, University of Calgary, 2500 University Drive NW, Calgary Alberta T2N 1N4, Canada; Department of Psychiatry, Cumming School of Medicine, University of Calgary, 28 Oki Dr, Calgary, AB T3B 6A8, Canada; Department of Educational Psychology, University of Alberta, 6-102 Education North, University of Alberta, Edmonton, AB T6G 2G5, Canada; Department of Pediatrics, Cumming School of Medicine, University of Calgary, 28 Oki Dr, Calgary, AB T3B 6A8, Canada; Cumming School of Medicine, University of Calgary, 3330 Hospital Dr NW, Calgary, AB T2N 4N1, Canada, IWK Health, 5980 University Ave #5850, Halifax, NS B3K 6R8; Department of Psychiatry, Dalhousie University, 5909 Veterans’ Memorial Lane, 8th Floor, Abbie J. Lane Memorial Building, QEII Health Sciences Centre, Halifax, NS B3H 2E2 Canada

**Keywords:** ADHD, executive functions, subcortical volumes, parent ratings

## Abstract

**Background:** The caudate and putamen have previously been implicated in Attention-Deficit/Hyperactivity Disorder (ADHD). However, previous studies have not investigated the relationship between the caudate and putamen with executive function (EF). The current study investigated the clinical relevance of the caudate and putamen with respect to EF.

**Method:** We studied 49 children (24 ADHD/25 typically developing children (TDC)). All participants in the ADHD group had to undergo a 48-hour stimulant medication washout period. Participants completed cognitive tasks related to working memory/inhibition and underwent a T1-weighted MRI sequence. All parents completed behaviour rating scales using the Behavior Rating Inventory of Executive Function, Second Edition (BRIEF-2). Data were analyzed using multivariate analysis of covariance, Pearson correlations, and linear regressions.

**Results:** Children with ADHD demonstrated a higher frequency of perseverative errors compared to TDC (*p <.*05), and their parents reported significantly more EF challenges *(p* <.001). No difference was observed in the working memory tasks. No significant volumetric differences were seen in the caudate or the putamen. A linear regression model suggested that the right caudate volume accounted for 10.3% of the variance in emotion regulation as reported by parents on the BRIEF-2 in the overall sample.

**Discussion:** We observed significant EF challenges without volumetric differences. However, the right caudate was correlated to parent ratings of emotional regulation, highlighting the need to consider emotional regulation difficulties in ADHD.

---

Subcortical regions, including the caudate and the putamen, have been previously implicated in Attention-Deficit/Hyperactivity Disorder (ADHD). ADHD is characterized by developmentally inappropriate levels of inattention or impulsivity and hyperactivity. Recent estimates suggest that the prevalence of ADHD ranges between 5-9% in Canadian school-aged children (Brault & Lacourse, 2012; Espinet et al., 2022; Polanczyk et al., 2014). Along with the primary symptoms, children diagnosed with ADHD struggle with their academic, motor, and social functioning (Barkley et al., 2001; Biederman et al., 2004; Wolraich et al., 2019), thereby increasing the use of societal resources such as increased annual medical costs compared to peers and reduced efficiency at work by parents (Hakkaart-van Roijen et al., 2007; Matza et al., 2005).

Children with ADHD often exhibit executive function (EF) challenges on working memory, response inhibition, set-shifting and planning tasks (Bünger et al., 2021; Kofler et al., 2019; Toplak et al., 2009; Willcutt et al., 2005). EF is an umbrella term used to describe higher-order goal-oriented processing skills (Diamond, 2013). These higher-order skills help with concentrating, paying attention, and controlling automatic behaviours; and are often associated with academic success (Borella et al., 2010; Cortés Pascual et al., 2019; Doebel, 2020), job success (Bailey, 2007; Chan et al., 2021) and overall quality of life (Dewey & Volkovinskaia, 2018; Schwörer et al., 2020; Stern et al., 2013). While some research studies have demonstrated that children with ADHD have EF deficits (Bünger et al., 2021; Kofler et al., 2011, 2019; Rapport et al., 2008; Willcutt et al., 2005), the findings are inconsistent with studies showing variable EF performance across the different domains likely impacted by a variety of factors including the difference between performance during testing compared to everyday functions, different conceptualizations of EF (no consensus regarding definitions), and sample characteristics (Huang-Pollock et al., 2012; Kofler et al., 2019; Willcutt et al., 2005). Estimates predict that between 21% to 60% of individuals with ADHD show impairment on EF tasks (Kofler et al., 2011). Also, concerns regarding construct validity and task impurity have been reported, with certain EF tasks requiring more than one set of executive and non-executive function skills (Kofler et al., 2019; Miyake & Friedman, 2012). Furthermore, comorbidities and intellectual functioning can also impact EF performance (Bental & Tirosh, 2007; Kofler et al., 2019).

In addition to EF challenges, children with ADHD exhibit neuroanatomical and neurochemical differences, particularly in brain regions that are related to EF skills (Altabella et al., 2014; Cortese et al., 2014; Cortese & Coghill, 2018; Hai et al., 2020; Hoogman et al., 2019; Kahl et al., 2022; MacMaster et al., 2003; Rubia, 2018). Studies have shown thinner prefrontal cortex in children with ADHD (Hai et al., 2022; Overmeyer et al., 2001; Shaw et al., 2007; Yang et al., 2015). A large-scale meta-analysis with over 2200 ADHD participants, including both pediatric and adult samples (mean age = 19.22 years, range = 4 – 62 years), found lower surface area, cortical thickness and reduced volumes in subcortical regions with more pronounced effect in children (Hoogman et al., 2017, 2019). However, baseline data from the Adolescent Brain and Cognitive Development Study cohort (ABCD) did not find any significant group difference in the caudate and putamen in 9-10 year olds (Bernanke et al., 2022). Moreover, differences in neurochemicals such as glutamate, N-acetyl aspartate, choline, and Gamma-aminobutyric acid (GABA) have been reported in children with ADHD compared to their age-matched peers (Edden et al., 2012; Hai et al., 2020; Kahl et al., 2022; MacMaster et al., 2003; Tafazoli et al., 2013). Overall, the current literature supports some neurochemical and neuroanatomical differences in ADHD.

While numerous studies have shown neuroanatomical and neurochemical differences, there is a paucity of studies to date that have combined neuroimaging, EF skills as reported by parents and EF task performance to understand better the neuroanatomy associated with EF skills (Almeida et al., 2010; Gau et al., 2015; Shang et al., 2013). It is essential to understand the relevance of structural differences with commonly used EF measures, as it will enable us to determine whether structural neuroimaging findings can be considered potential biomarkers of ADHD.

## Research Purpose

The primary purpose of this study is to investigate EF differences in children with ADHD compared to typically developing controls (TDC) using performance-based tasks (working memory and inhibition tasks) and behaviour ratings completed by parents. The secondary purpose of this study is to examine differences in the volume of the subcortical regions (caudate and putamen) of children with ADHD compared to TDC. Importantly, this study will take on a multimodal approach to understand processes impacted in children with ADHD.

## Methods

The current study recruited a total of 55 participants. Two participants were excluded because they did not meet the study eligibility criteria: one participant had a diagnosis of ASD, and one participant failed to observe the 48-hour medication washout period. Also, one participant withdrew within one hour of joining due to extreme shyness and anxiety. Lastly, three additional participants were excluded due to their subcortical volumes data being outliers, as indicated through quality control measures. A final sample of 49 (24 children with ADHD (age range 7.53-16.4 years old, *SD* = 2.63) and 25 TDC (age range 7.26-16.87 years, *SD* = 2.76)) between the ages of 7–16 years old were included in the present analyses. All participants completed EF assessments comprising inhibition and working memory. All participants also underwent T1-weighted structural MRI. The study received research ethics approval from the Conjoint Health Research Ethics Board (CHREB) at the University of Calgary (REB19-0499).

### Neuropsychological Measures

This study measured two primary domains of EF (working memory and response inhibition) in both children with ADHD and a typically developing control group. These two EF domains were chosen as these are the most commonly studied EF skills in the developmental literature, with empirical support suggesting limited overlap with other EF skills (McAuley & White, 2011). The Digit Span Backwards and Spatial Span Backwards subtests were used to measure working memory from the Wechsler Intelligence Scale for Children (WISC-V) and WISC-V Integrated (Wechsler, 2014). Commission, Omission and Perseverative errors on the Conners’ Continuous Performance Test (CPT III; (K. Conners, 2014) and Colour Word Interference test from the Delis Kaplan Executive Function System (D-KEFS) were used to measure inhibition (Delis et al., 2001).

### Behavioural Measures-Parent Rating Scale

Parents completed two behaviour rating scales for the current study: i) Conners-3 Parent Rating Scale (Conners, Keith, 2008) and ii) Behavior Rating Inventory of Executive Function, Second Edition (BRIEF-2; (Gioia et al., 2015). Conners-3 rating scales have high levels of internal consistency, with Cronbach’s alpha ranging from 0.77 to 0.97 (mean Cronbach’s alpha = 0.90), and test-retest correlations ranging from 0.71 to 0.98 (C. K. Conners et al., 2011). The BRIEF-2 exhibited excellent internal consistency (Cronbach’s α = 0.97). Cronbach’s alpha scores for the different subscales ranged from .77 to .92 (Jacobson et al., 2016).

#### MRI Acquisition Protocol

All participants underwent a high-resolution MRI T1-weighted sequence using a 3 Tesla General Electric Discovery 750W MRI scanner with a 32-channel head coil. Structural MRI parameters were as follows: TR = 8.2ms, TE = 3.2ms, flip angle = 10°, field of view (FOV) = 256 mm^2^, acquisition matrix size = 300×300, voxels = 0.8mm^3^ isotropic.

### Data Collection Procedures

Participants were recruited through flyers posted around the university campus, referrals from healthcare professionals in the general community, and social media such as Facebook and Twitter. All participants in the ADHD group had to have a prior diagnosis of ADHD reported by a physician or a psychologist. First, informed consent from the parents and assent from participants was obtained. Then, parents and the participants completed the MINI-KID, a semi-structured interview to evaluate eligibility for the study (Sheehan et al., 2010). Parents also completed Conners-3 rating scales to confirm their child’s ADHD diagnosis. Following receiving informed consent from parents and assent from the children, the author also completed the intellectual Quotient (IQ) screener that included the three subtests from the WISC-V and WISC-V Integrated with the participants.

All eligible participants then completed the additional neuropsychological measures, with parents completing the questionnaires in one assessment room and the child completing the assessments in a separate assessment room. The testing session lasted approximately 90 minutes. After completing the tasks, eligible participants underwent the neuroimaging portion which took one hour and fifteen minutes to complete. All participants received a gift card valued at $150.00 CDN for their participation in the research study.

#### MRI Reconstruction of Subcortical Regions

FreeSurfer Software (Fischl & Dale, 2000) was used to obtain the subcortical volumes (See Figure 1 and 2). A detailed description of the FreeSurfer processing is described online (https://surfer.nmr.mgh.harvard.edu/fswiki/FreeSurferMethodsCitation; (Dale et al., 1999; Fischl et al., 2002, 2004; Fischl & Dale, 2000). This measurement technique has been validated in both adult and pediatric populations (Biffen et al., 2020; J. Dewey et al., 2010).

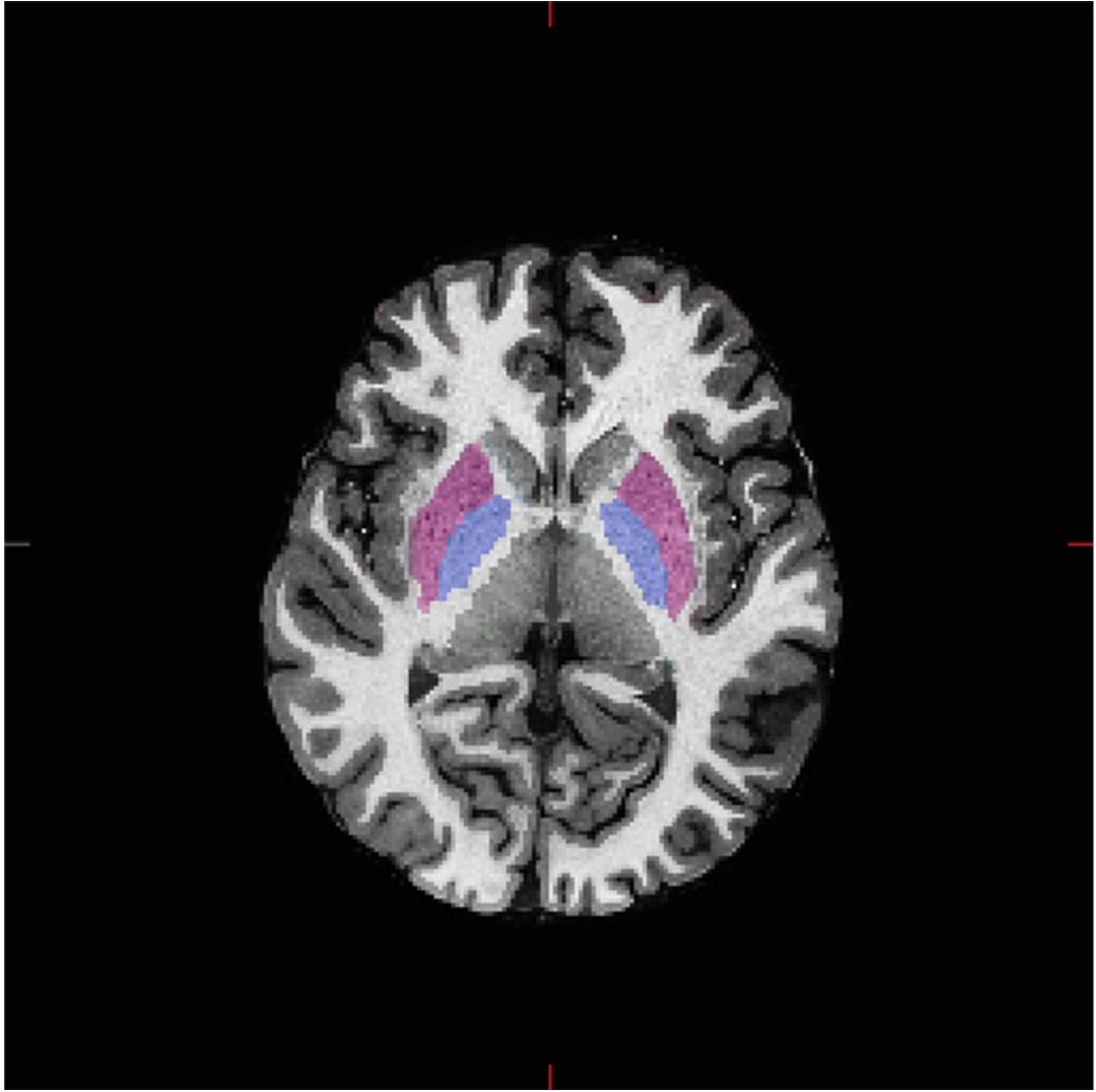

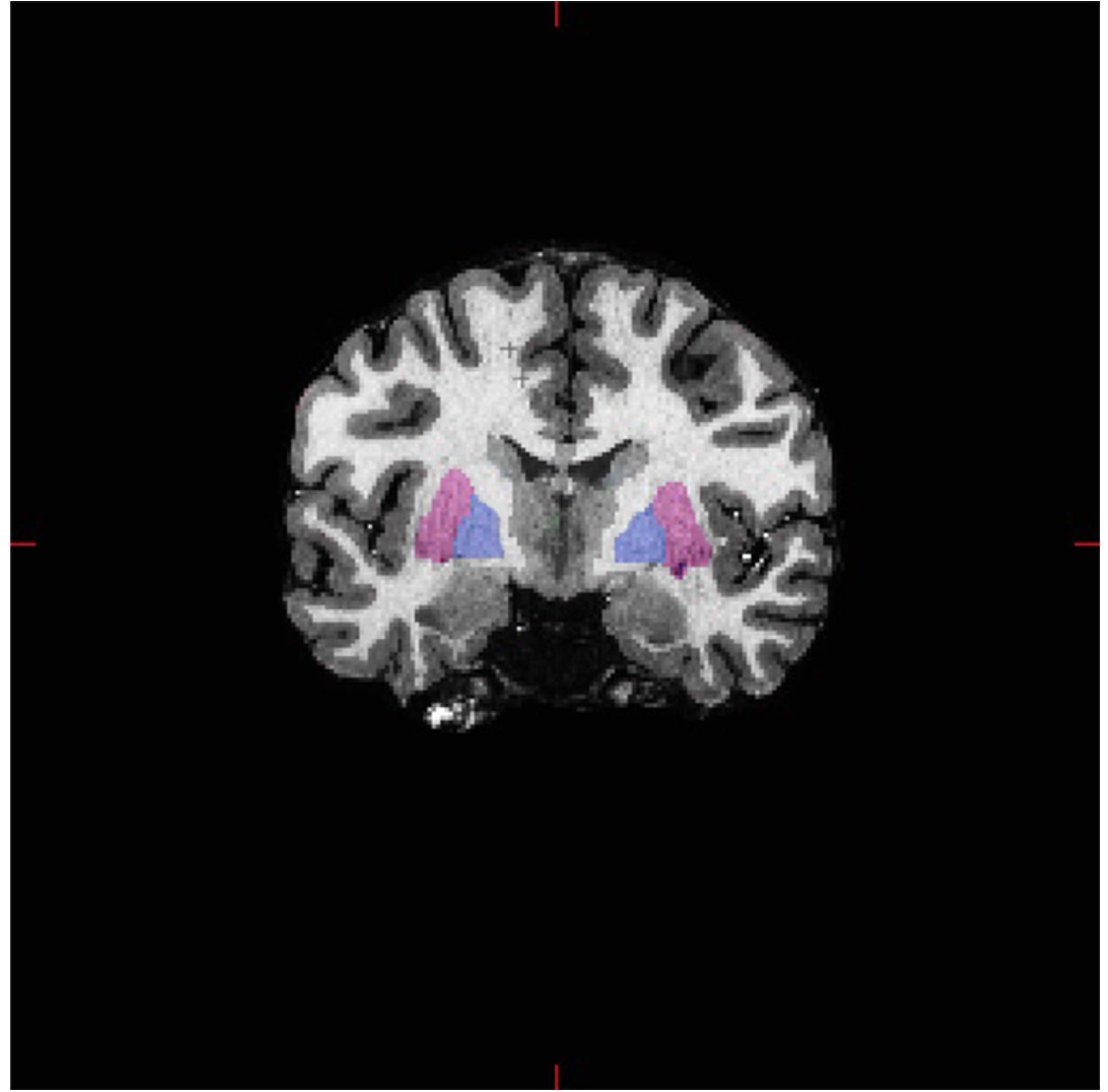

### Data Analysis Plan

SPSS version 29.0 was used to conduct all the planned data analyses. Multivariate analyses of variance (MANOVA) were conducted to investigate the differences in performance on the EF tasks and BRIEF-2 parent measures between children with ADHD and the TDC group. Benjamini-Hochbergs Principle was used to correct for multiple comparisons. Multivariate Analysis of Covariance (MANCOVA) was carried out to investigate the differences in volumes of the caudate and putamen in children with ADHD compared to the TDC group, with total intracranial volume (ICV), age and biological sex as covariates. Additional post-hoc analysis was conducted to investigate group differences between the bilateral amygdala and hippocampus. Pearson and Spearman correlations were conducted to investigate the associations between the caudate and putamen volumes with EF skills and linear regressions were conducted using variables with significant correlations.

## Results

The data were inspected for missing values and outliers prior to running any statistical analyses. The data were also evaluated for normality, linearity, homogeneity of variance, and homoscedasticity to meet the assumptions of the parametric analysis.

### Group Differences in Screening Measures

There was no age, biological sex or any other sociodemographic factors differences between the two groups (see Table 1 for demographic information and baseline information). As expected, there were significant group differences in ADHD symptoms, as reported by parents on the Conners-3 rating scale. Specifically, parents of children with ADHD endorsed higher levels of Inattentive (*t* (47) = 7.08, *p* < .001, Cohen’s *d* = 2.02) and Hyperactive/Impulsivity (*t* (47) = 8.83, *p* < .001, Cohen’s *d* = 2.51) symptoms compared to the TDC group. There was no significant group difference in the three intellectual functioning screener that were completed: WISC-V Arithmetic subtest (*t* (47) = 1.57, *p* = .12, Cohen’s *d* = 0.45), WISC-V Integrated Vocabulary subtest (*t* (47) = .90, *p* = .37, Cohen’s *d* = 0.26) and WISC-V Integrated Block Design subtest (*t* (47) = 0.32, *p* = .75, Cohen’s *d* = 0.09).

**Table 1.**
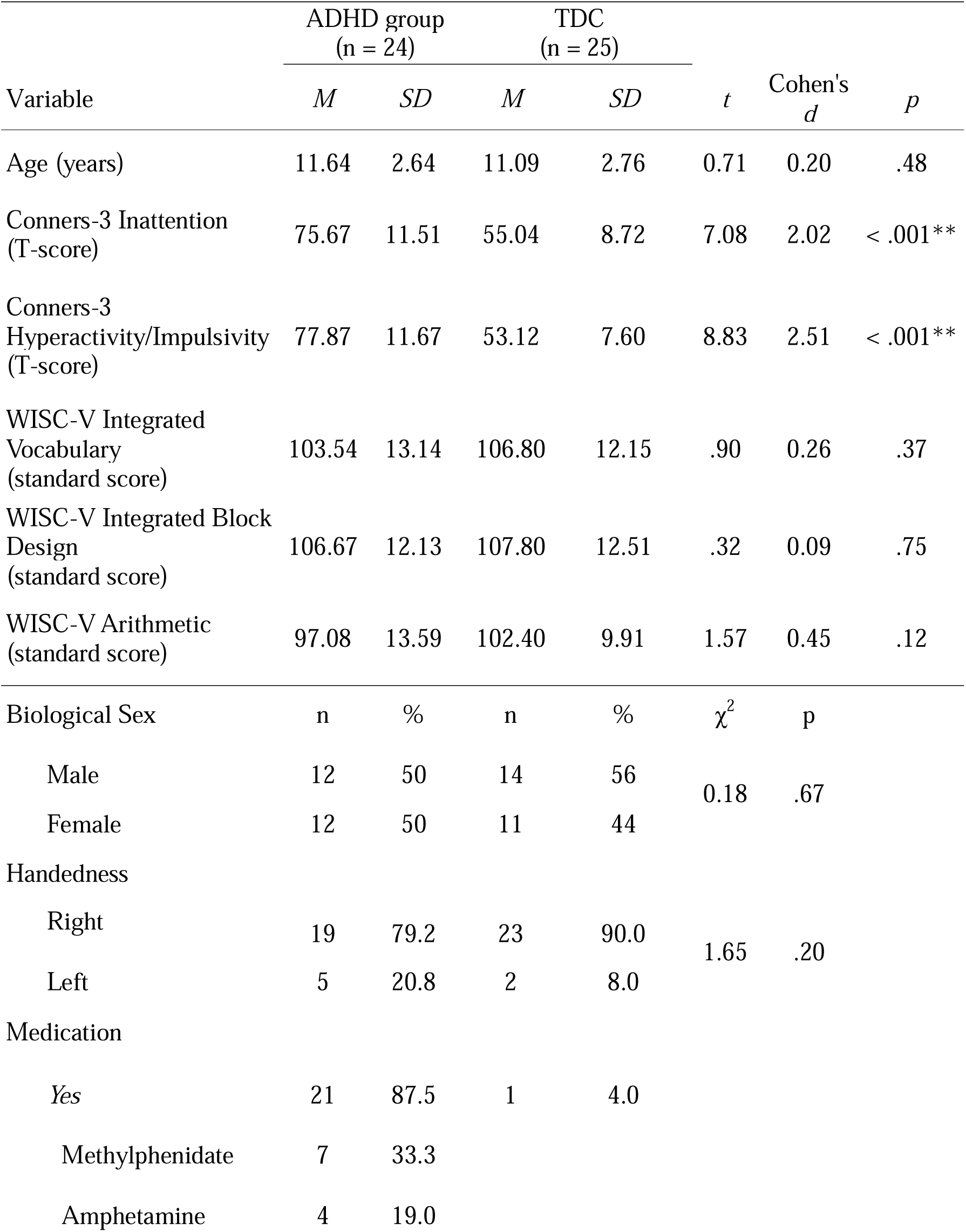
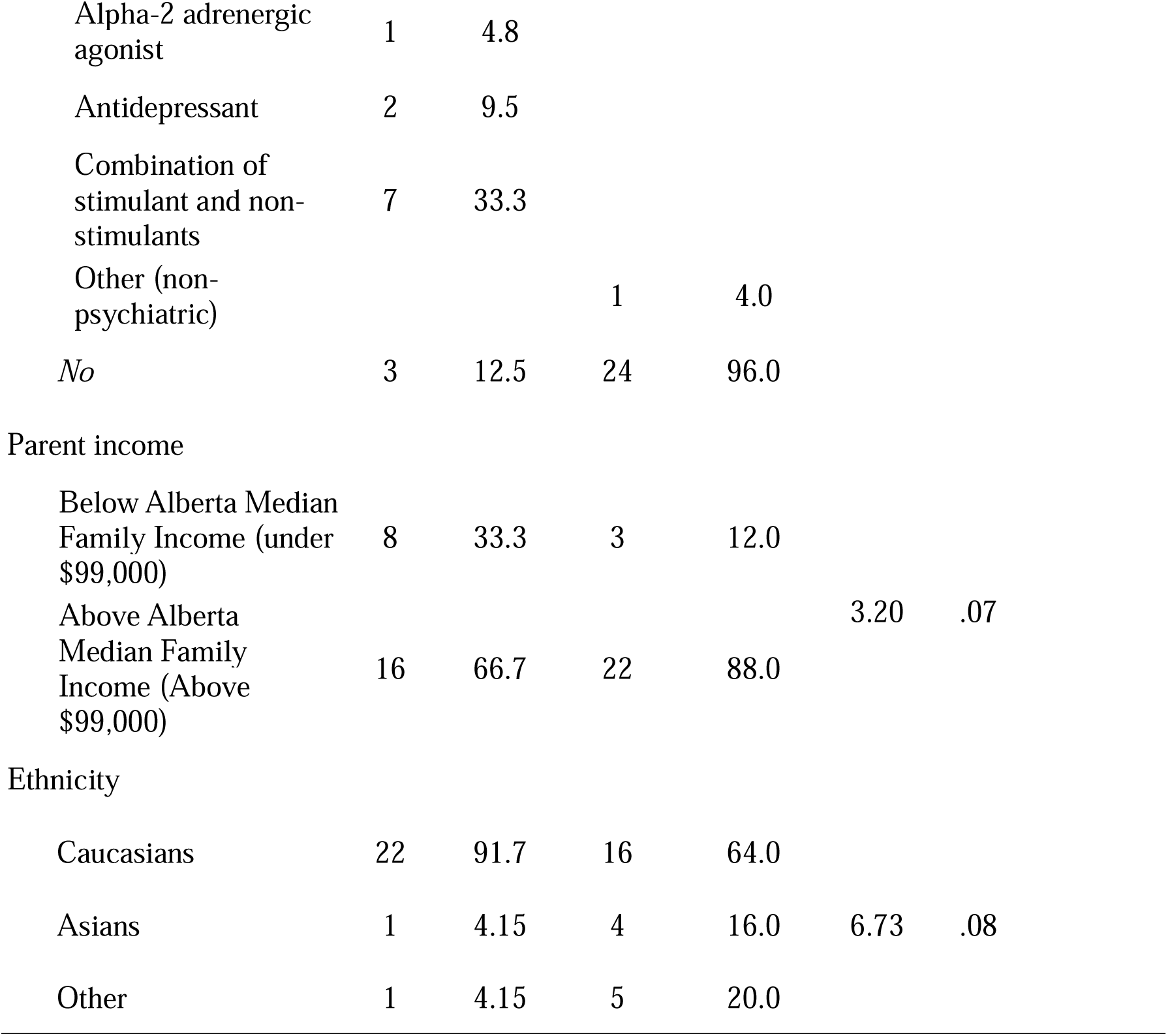
Participant characteristic information, including demographic information, intellectual functioning test results, and ADHD symptoms.

| Variable | ADHD group<br>(n = 24) |  | TDC<br>(n = 25) |  | *t* | Cohen's<br>*d* | *p* |
| --- | --- | --- | --- | --- | --- | --- | --- |
|  | *M* | *SD* | *M* | *SD* |  |  |  |
| Age (years) | 11.64 | 2.64 | 11.09 | 2.76 | 0.71 | 0.20 | .48 |
| Conners-3 Inattention<br>(T-score) | 75.67 | 11.51 | 55.04 | 8.72 | 7.08 | 2.02 | < .001** |
| Conners-3<br>Hyperactivity/Impulsivity<br>(T-score) | 77.87 | 11.67 | 53.12 | 7.60 | 8.83 | 2.51 | < .001** |
| WISC-V Integrated<br>Vocabulary<br>(standard score) | 103.54 | 13.14 | 106.80 | 12.15 | .90 | 0.26 | .37 |
| WISC-V Integrated Block<br>Design<br>(standard score) | 106.67 | 12.13 | 107.80 | 12.51 | .32 | 0.09 | .75 |
| WISC-V Arithmetic<br>(standard score) | 97.08 | 13.59 | 102.40 | 9.91 | 1.57 | 0.45 | .12 |
| Biological Sex | n | % | n | % | $\chi^2$ | p | |
| Male | 12 | 50 | 14 | 56 | 0.18 | .67 |  |
| Female | 12 | 50 | 11 | 44 |  |  |  |
| Handedness |  |  |  |  |  |  |  |
| Right | 19 | 79.2 | 23 | 90.0 | 1.65 | .20 |  |
| Left | 5 | 20.8 | 2 | 8.0 |  |  |  |
| Medication |  |  |  |  |  |  |  |
| Yes | 21 | 87.5 | 1 | 4.0 |  |  |  |
| Methylphenidate | 7 | 33.3 |  |  |  |  |  |
| Amphetamine | 4 | 19.0 |  |  |  |  |  |

|  |  |  |  |  |  |  |
| --- | --- | --- | --- | --- | --- | --- |
| Alpha-2 adrenergic agonist | 1 | 4.8 |  |  |  |  |
| Antidepressant | 2 | 9.5 |  |  |  |  |
| Combination of stimulant and non-stimulants | 7 | 33.3 |  |  |  |  |
| Other (non-psychiatric) |  |  | 1 | 4.0 |  |  |
| *No* | 3 | 12.5 | 24 | 96.0 |  |  |
| Parent income |  |  |  |  |  |  |
| Below Alberta Median Family Income (under \$99,000) | 8 | 33.3 | 3 | 12.0 | | |
| Above Alberta Median Family Income (Above \$99,000) | 16 | 66.7 | 22 | 88.0 | 3.20 | .07 |
| Ethnicity |  |  |  |  |  |  |
| Caucasians | 22 | 91.7 | 16 | 64.0 |  |  |
| Asians | 1 | 4.15 | 4 | 16.0 | 6.73 | .08 |
| Other | 1 | 4.15 | 5 | 20.0 |  |  |

### EF differences based on Performance-Based Tasks

MANOVA did not show any group differences on the working memory task performance between the ADHD and TDC groups (*F* (2, 46) = 1.38, *p* =.26, partial eta squared = .06). Similarly, no group differences on the response inhibition task performance between the ADHD and TDC groups were observed (*F* (4, 39) = 2.48, *p* =.06, partial eta squared = .20). However, univariate analysis of variance showed that children with ADHD made more perseverative errors than the TDC group on the CPT-3 task (*F* (1, 42) = 8.18, *p* = .007, partial eta squared = .16), with Benjamini-Hochbergs corrected *p* = .04. No other significant differences in performance were observed (see Table 2 for more details).

**Table 2.** Executive Function Performance Scores of the ADHD and Typically Developing Control (TDC) Groups.

| Variable | ADHD |  | TDC |  | MANOVA |  |  |
| --- | --- | --- | --- | --- | --- | --- | --- |
|  | (n = 24) |  | (n = 25) |  |  |  |  |
|  | *M* | *SD* | *M* | *SD* | *F* | *p* | Partial Eta squared |
| *Working Memory* |  |  |  |  |  |  |  |
| Digit Span Backwards (standard score) | 96.88 | 12.49 | 102.40 | 11.28 | 2.64 | .11 | .05 |
| Spatial Span Backwards (standard score) | 101.46 | 15.98 | 101.40 | 13.11 | .00 | .99 | .00 |
| *Response Inhibition* |  |  |  |  |  |  |  |
| CPT-3 Omission Errors T-score (n=22) | 62.41 | 16.72 | 54.61 | 13.99 | 2.67 | .11 | .06 |
| CPT-3 Commission Errors T-score (n=22) | 56.86 | 7.58 | 55.18 | 7.81 | .53 | .47 | .01 |
| CPT-3 Perseverative Errors T-score (n=22) | 68.77 | 17.00 | 54.95 | 14.97 | 8.18 | .007 | .16** |
| DKEFS Inhibition (standard score) | 98.41 | 14.09 | 103.18 | 11.81 | 1.48 | .23 | .03 |

\*\* Indicates *p* value < .01

### EF differences based on Parent Ratings

The EF results are summarized in Table 3. The results from the MANOVA indicated significant difference in parent ratings between children with ADHD and the TDC group (*F* (5,43) = 20.89, *p* <.001, partial eta squared =.71). Specifically, across all the index scores, parents of children with ADHD reported significantly higher EF challenges compared to the TDC. These increased EF challenges were reported on all three subscales, Behaviour Regulation Index (*F* (1,47) = 53.44, *p* <.001, partial eta square =.53), Emotion Regulation Index *F* (1,47) = 31.26, *p* <.001, partial eta square =.40, and Cognitive Regulation Index *F* (1,47) = 64.86, *p* <.001, partial eta square =.58. Furthermore, significant challenges were also reported by parents on scales specifically related to Inhibition, *F* (1,47) = 46.48, *p* <.001, partial eta square =.50 and Working Memory *F* (1,47) = 106.88, *p* <.001, partial eta square =.70.

**Table 3.**
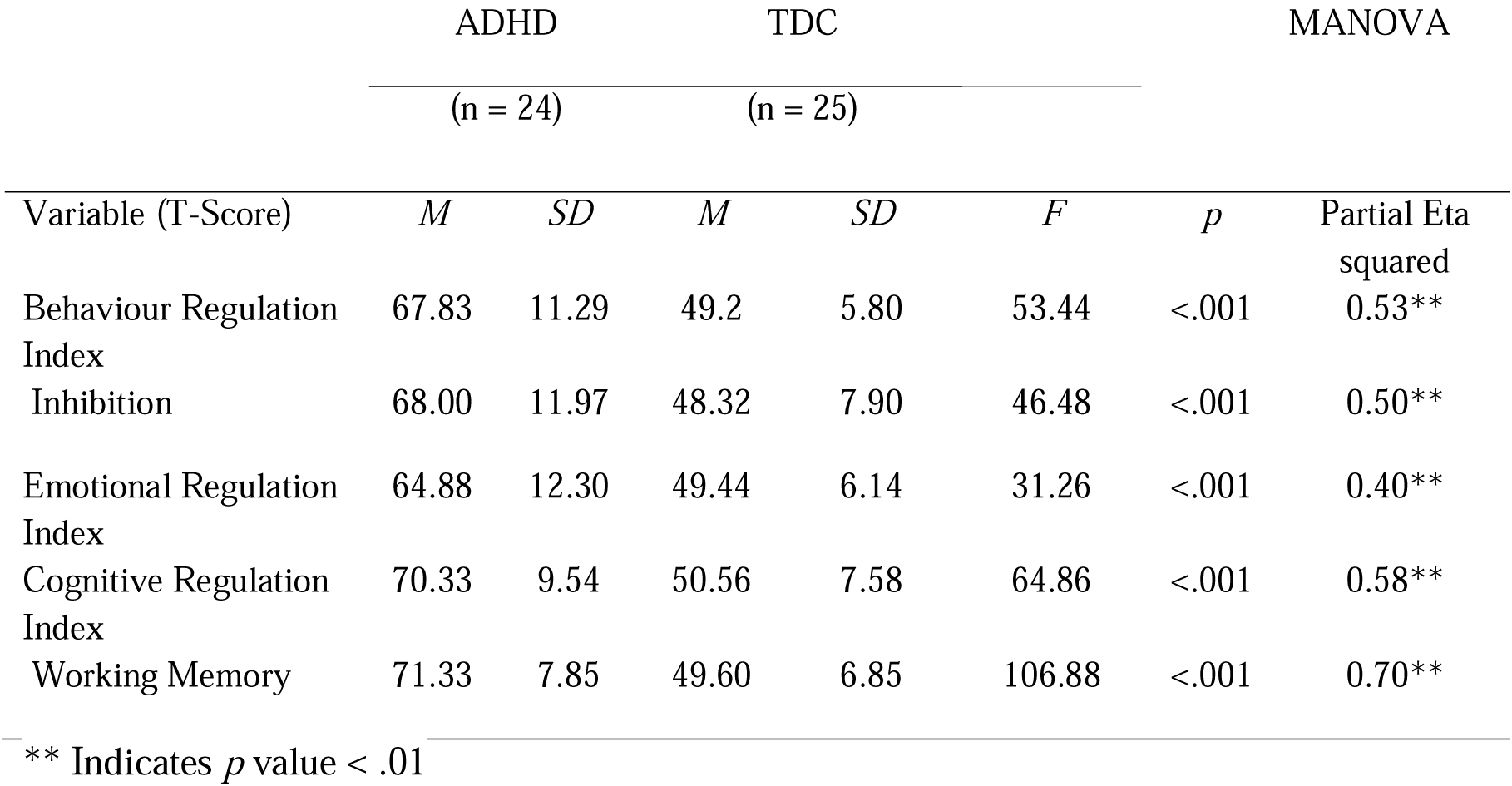
Parent Ratings of Executive Function (BRIEF-2) in the ADHD and Typically Developing Control (TDC) Groups.

|  | ADHD |  | TDC |  | MANOVA |  |  |
| --- | --- | --- | --- | --- | --- | --- | --- |
|  | (n = 24) |  | (n = 25) |  |  |  |  |
| Variable (T-Score) | *M* | *SD* | *M* | *SD* | *F* | *p* | Partial Eta squared |
| Behaviour Regulation Index | 67.83 | 11.29 | 49.2 | 5.80 | 53.44 | <.001 | 0.53** |
| Inhibition | 68.00 | 11.97 | 48.32 | 7.90 | 46.48 | <.001 | 0.50** |
| Emotional Regulation Index | 64.88 | 12.30 | 49.44 | 6.14 | 31.26 | <.001 | 0.40** |
| Cognitive Regulation Index | 70.33 | 9.54 | 50.56 | 7.58 | 64.86 | <.001 | 0.58** |
| Working Memory | 71.33 | 7.85 | 49.60 | 6.85 | 106.88 | <.001 | 0.70** |

\*\* Indicates *p* value < .01

### Subcortical Differences

Table 4 summarizes the subcortical findings. The results from the MANCOVA showed no significant group differences in the volumes of the right and left caudate and putamen, *F* (4,41) = .79, *p* >.05, partial eta square =.07). Additional post-hoc analyses were conducted to identify group differences in the bilateral hippocampus and amygdala. Significant group difference was observed in the left amygdala only (*F* (1, 46) = .4.48, *p* <.05, partial eta square =.07). However, this group difference did not survive multiple comparison tests using Benjamini-Hochberg method. No other group differences were observed.

**Table 4.**
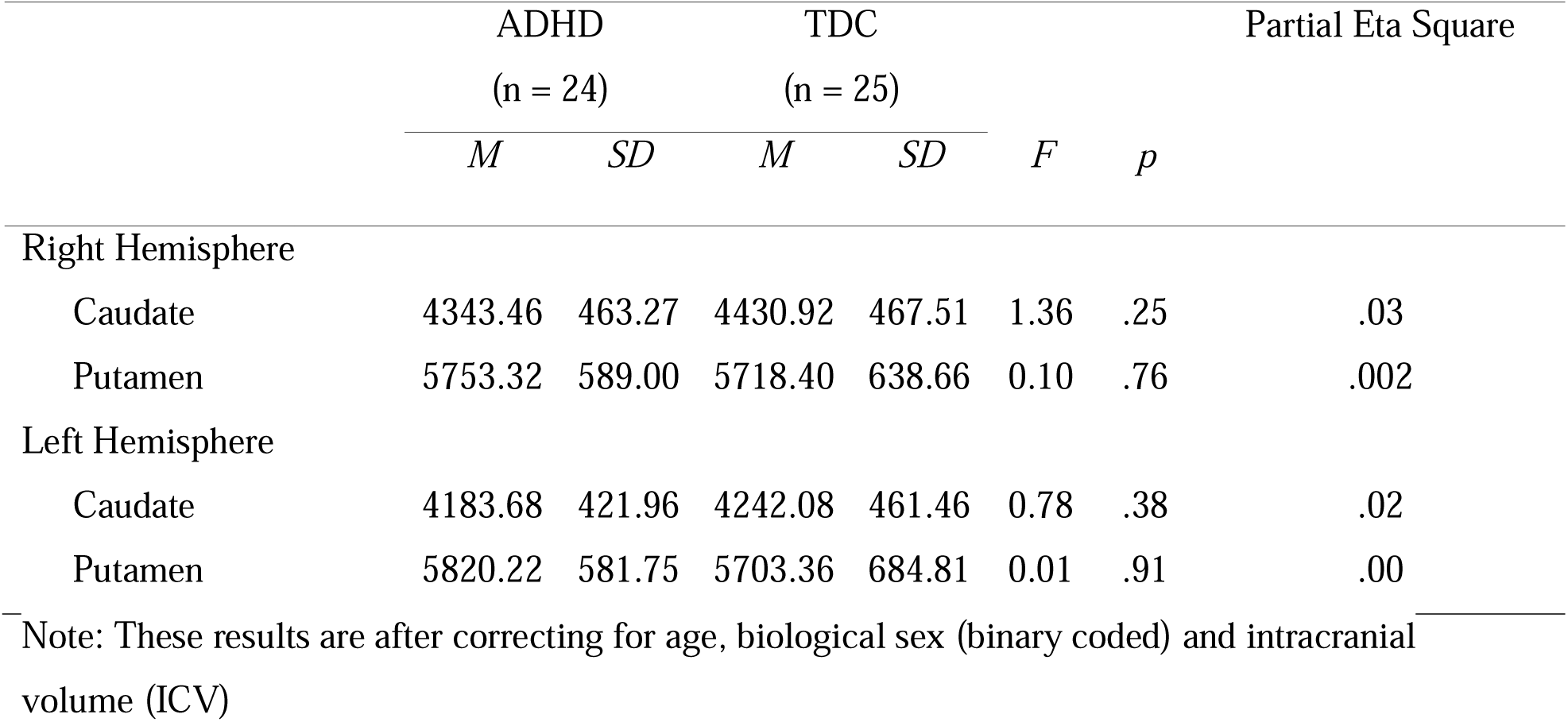
Subcortical Volume Measurements of the ADHD and Typically Developing Control (TDC) Groups.

### Brain behaviour Relationships

No significant correlations were observed between caudate and putamen volumes with any of the performance-based EF measures. Significant correlations were observed with right caudate volume and parent ratings on the BRIEF-2 ERI subscale (*r* = -.32, *p* = .02). No other significant correlations were found with any of the other BRIEF-2 subscales with the right caudate. Similarly, no other significant correlations were observed with the BRIEF-2 subscales with left caudate, right putamen and left putamen volumes.

When correlations were conducted separately for the ADHD and TDC group, significant correlations were observed for the ADHD group for right caudate volume with BRIEF-2 ERI subscale (*r* = -.51, *p* = .01; See Table 5 for more details).). No such correlation was observed in the TDC group on the BRIEF-2 ERI subscale (*r* = -.06, *p* = .77).

**Table 5.** Correlations between Right Caudate Volume and Parent Ratings of EF Skills (BRIEF-2) in the ADHD group.

|  | Variables | 1 | 2 | 3 | 4 | 5 | 6 |
| --- | --- | --- | --- | --- | --- | --- | --- |
| 1 | Right Caudate | - | $r=-.251$<br>$p=.237$ | $r=-.513^*$<br>$p=.010$ | $r=-.067$<br>$p=.757$ | $r=-.286$<br>$p=.175$ | $r=-.105$<br>$p=.625$ |
| 2 | BRIEF-2 Behaviour Regulation Index | | - | $r=.599^{**}$<br>$p=.002$ | $r=.653^{**}$<br>$p=.001$ | $r=.953^{**}$<br>$p=.000$ | $r=.601^{**}$<br>$p=.002$ |
| 3 | BRIEF-2 Emotional Regulation Index | | | - | $r=.507^*$<br>$p=.011$ | $r=.596^{**}$<br>$p=.002$ | $r=.525^{**}$<br>$p=.009$ |
| 4 | BRIEF-2 Cognitive Regulation Index | | | | - | $r=.617^{**}$<br>$p=.001$ | $r=.881^{**}$<br>$p=.000$ |
| 5 | BRIEF-2 Inhibition | | | | | - | $r=.588^{**}$<br>$p=.003$ |
| 6 | BRIEF-2 Working Memory |  |  |  |  |  | - |

\*\* indicates correlation significant at the 0.01 level (2-tailed).
\* Indicates correlation is significant at the 0.05 level (2-tailed).

Linear Regressions were completed with BRIEF-2 ERI subscale and right caudate volume for the total sample as well as with the ADHD group only. Results showed that 10.3% of the variance in the BRIEF-2 ERI subscale was explained by the right caudate volume for the overall sample, (*F* (1, 47) = 5.41, *p* =. 02, *R^2^* = 0.10. Similarly, 26.3 % of the variance in the BRIEF-2 ERI subscale was explained by the right caudate volume for the ADHD group only, (*F* (1, 22) = 7.85, *p* = 0.01, *R^2^* = 0.263.

## Discussion

This study demonstrated that there are EF challenges observed in children with ADHD compared to their typically developing peers. These differences were reported by parents on the behaviour rating scale. On performance-based measures of EF, the difference was less pronounced, with children with ADHD making more perseverative errors on the response inhibition task compared to their peers. This alludes to the importance of using multimodal measures to better understand EF challenges in ADHD. Regarding subcortical volumes, no significant group differences were observed in either the caudate or putamen of children with ADHD compared to TDC. However, the current study showed some exploratory findings that could possibly indicate the clinical relevance of the right caudate volume in relation to emotional regulation challenges observed in children with ADHD, as reported by parents. Specifically, this correlation was observed in the full study sample, while driven primarily by the ADHD group. Thus, suggestive of the caudate and its involvement in emotional regulation in children with ADHD.

EF difficulties are often reported in children with ADHD based on objective measures (Kofler et al., 2019; Willcutt et al., 2005). However, our study did not show any overall significant group differences in EF based on performance on objective cognitive tasks. The only significant difference was observed on a response inhibition task, where children with ADHD made more perseverative errors than their peers. This group difference was significant, following controlling for multiple comparisons. No other significant group difference was observed on any of the other measures of response inhibition or spatial and auditory working memory. This inconsistency in EF findings align with the existing literature as some studies have reported neurocognitive heterogeneity in children with ADHD as a common phenomenon (Harmon et al., 2018; Kofler et al., 2019; Nigg et al., 2005). This heterogeneity in presentation could be for numerous reasons. For example, Kofler et al. (2019) reported that only 35% of children with ADHD were impaired when EF was considered a unitary construct. However, this prevalence rate changed to 89% when EF impairment was considered multi-dimensional and observed in at least one of the three primary EF domains (working memory, inhibition or set-shifting). Our results support of the importance of using a variety of EF measures to fully understand the impact of ADHD with recommendations for best clinical practice to include multiple EF domains in the assessment.

While performance tasks did not show significant EF difficulties, parents of children with ADHD reported challenges on all three subscales of the BRIEF-2, Behavioural Regulation Index (BRI), Emotional Regulation Index (ERI) and Cognitive Regulation Index (CRI), in children with ADHD compared to their typically developing peers. These findings are consistent with the existing literature where parents of children with ADHD frequently report struggles on the behaviour rating scales (Schneider et al., 2020; Toplak et al., 2009). These EF difficulties stated by parents represent a more global perspective and require parents to estimate EF challenges over the past six months. Furthermore, these findings suggest that children with ADHD not only struggle with common EF domains such as working memory and inhibition but in other areas as well. For example, parents reported observing difficulties in skills such as planning, task monitoring, and initiating. Additionally, it is important to consider the impact of parents quality of life, parental stress and challenges associated with parenting children with ADHD and the influence on rating scales (Joyner et al., 2009).

One of the interesting findings from the current study is the challenges reported by parents around emotional regulation. Emotional regulation challenges are quite prevalent in children with ADHD and were included as a diagnostic criterion (Faraone et al., 2019; Shaw et al., 2014). This conceptualization was seen in the earlier diagnostic criteria for ADHD but was changed in the DSM-III (APA, 2022). However, children with ADHD often show more negative affect, temper outbursts, and emotional lability when presented with negative and challenging situations (Rohr et al., 2021; Shaw et al., 2014). While emotional dysregulation is not uncommon in other disorders, children diagnosed with ADHD frequently show emotional dysregulation in the form of reduced patience, unwillingness to wait their turn, high frustration, anger, and irritability (Barkley & Fischer, 2010; Faraone et al., 2019). It has also been suggested that children with ADHD often struggle with downregulating these high-intensity emotions as they lack the ability to use their EF skills to regulate their emotions and behaviour.

The third research question of the current study was to investigate volumetric differences in subcortical regions in children with ADHD compared to their peers. The results found no significant volumetric differences between children with ADHD and their peers in either the caudate or the putamen after controlling for age, biological sex and ICV. These findings are in contrast to the expected hypotheses, as previous studies, including a large-scale meta-analysis, showed altered volumes in these subcortical regions, observed in children, adolescents and adults, with larger effect sizes observed in children (Hoogman et al., 2017). However, the age ranges included in the stratified analysis were variable, with participants younger than 15 years old considered to be in the children group, adolescents aged 15–21 years, and adults aged 22 years and older. Furthermore, the participants in the present study are well characterized with extensive behavioural, cognitive, and neuroanatomical information, which provides important clinically relevant information about the sample.

Despite these prior findings on subcortical structures, the current study’s results are not unexpected, given the inconsistency in existing literature regarding neuroanatomical differences observed in children with ADHD. Recent studies, notably a large-scale study from the ABCD cohort by Bernanke et al. (2022) and others, have also reported no significant group differences in the subcortical regions, contributing to the ongoing debate surrounding this issue (Bernanke et al., 2022; Cortese & Coghill, 2018; Rubia, 2018; Samea et al., 2019). One reason for the differences could be the use of medication (duration and type of medications) and its impact on different brain regions. The children who took part in the current study were not medication naïve. While they had undergone a 48-hour washout period, it is difficult to rule out the impact of medication use long-term. In addition, medication compliance can also impact the efficacy of medication use and the overall long-term impact on different brain regions. Furthermore, some children with ADHD often take “holiday” breaks from their medication, which also makes it challenging to statistically control for the impact of long-term medication use and its effects on different brain regions.

The novelty of the current study is the significant correlations between right caudate volume and EF parent rating scales related to emotional regulation. The relationship indicated a negative correlation where a higher volume of right caudate was generally associated with lower emotional regulation challenges in children with ADHD. The regression model further showed that right caudate volume predicted 10.3 % of the variance in BRIEF-2 ERI scores in the total sample. Overall, it suggests a possible brain behaviour relation between right caudate volume and emotion regulation challenges. While these results are exploratory, it is possible that the caudate may serve as a link between the basal ganglia, dopaminergic reward systems, and ADHD symptomatology (Damiani et al., 2021). Moreover, given that the cortico-striatal network is associated with behavioural features characterizing ADHD —such as difficulty tolerating delays and impulsivity—it is possible that challenges in emotional regulation may be tied to the manifestation of ADHD symptoms.

### Limitations and Future Studies

While the current study adds valuable information, the results should still be evaluated in the context of some study limitations. First, the ADHD sample in the current study was based on self-referrals and it is possible that families who were involved in the study included a unique subset of the population. Given the time commitment required for the study participation (8+ hours), the study may include participants who are generally managing their ADHD symptoms well. Also, with improvement in MR technology future research needs to investigate larger cohorts of ADHD individuals to investigate other potential subgroups of ADHD based on neuroanatomical differences to offer more targeted interventions.

Further research is needed to explore the brain-behaviour relationships using different versions of the BRIEF-2, such as those completed by teachers, adolescents, and preschoolers. These rating scales, when used at various time points, can provide insights into the trajectory of brain development in children. Future research should also consider incorporating other EF tasks that are dependent on emotional regulation to enhance our understanding of the relationship between parent ratings of emotion regulation and performance on emotion regulation tasks. Finally, this study only assess EF and subcortical volumes as they pertain to ADHD. In the future, to better understand the etiology of ADHD, the impact of different factors such as parent-child relationships, environmental factors, school experience, and peers on brain development needs to be studied (Doom et al., 2021). Other biological factors such as hormonal change, impact of puberty, sleep, diet, and physical activity need to be incorporated in future models.

### Conclusion

Overall, this study showed significant EF difficulties based on parent ratings, but no statistically significant volumetric difference was observed in the caudate or putamen. However, the right caudate was correlated to parent ratings of emotional regulation and predicted 10.3 % of the variability in parent ratings. These findings highlight the need to consider emotional regulation difficulties in ADHD not just for diagnostic purposes but also for targeted treatment options. The current study also recommends assessing EF using a variety of assessment tools, such as informant rating scales and neuropsychological measures, for best practice. Given the exploratory nature of the findings, future studies are required to replicate current results and determine the role of caudate in emotional regulation.

## Acknowledgement

This work was supported by funding from the Alberta Children’s Hospital Foundation, the Werklund School of Education (University of Calgary), and the Alberta Graduate Excellence Scholarship awarded to TH. The authors would like to thank the families who took part in the study, as well as Hanna Frank and Kristina Lynberg for their support with data collection.

## Research Ethics

The study received research ethics approval from the Conjoint Health Research Ethics Board (CHREB) at the University of Calgary (REB19-0499). All parents provided consent for participation and the children provided assent.

## References

Almeida, L. G., Ricardo-Garcell, J., Prado, H., Barajas, L., Fernández-Bouzas, A., Ávila, D., & Martínez, R. B. (2010). Reduced right frontal cortical thickness in children, adolescents and adults with ADHD and its correlation to clinical variables: A cross-sectional study. Journal of Psychiatric Research, 44(16), 1214–1223. 10.1016/j.jpsychires.2010.04.026

Altabella, L., Zoratto, F., Adriani, W., & Canese, R. (2014). MR imaging–detectable metabolic alterations in attention deficit/hyperactivity disorder: From preclinical to clinical studies. American Journal of Neuroradiology, 35(6 suppl), S55 LP–S63. 10.3174/ajnr.A3843

Bailey, C. E. (2007). Cognitive Accuracy and Intelligent Executive Function in the Brain and in Business. Annals of the New York Academy of Sciences, 1118(1), 122–141. 10.1196/annals.1412.011

Barkley, R. A., Edwards, G., Laneri, M., Fletcher, K., & Metevia, L. (2001). Executive Functioning, Temporal Discounting, and Sense of Time in Adolescents With Attention. Journal of Abnormal Child Psychology, 29(6), 541–556.

Barkley, R. A., & Fischer, M. (2010). The unique contribution of emotional impulsiveness to impairment in major life activities in hyperactive children as adults. Journal of the American Academy of Child and Adolescent Psychiatry, 49(5), 503–513. 10.1097/00004583-201005000-00011

Bental, B., & Tirosh, E. (2007). The relationship between attention, executive functions and reading domain abilities in attention deficit hyperactivity disorder and reading disorder: A comparative study. Journal of Child Psychology and Psychiatry, 48(5), 455–463. 10.1111/j.1469-7610.2006.01710.x

Bernanke, J., Luna, A., Chang, L., Bruno, E., Dworkin, J., & Posner, J. (2022). Structural brain measures among children with and without ADHD in the Adolescent Brain and Cognitive Development Study cohort: A cross-sectional US population-based study. The Lancet Psychiatry, 9(3), 222–231. 10.1016/S2215-0366(21)00505-8

Biederman, J., Monuteaux, M. C., Doyle, A. E., Seidman, L. J., Wilens, T. E., Ferrero, F., Morgan, C. L., & Faraone, S. V. (2004). Impact of executive function deficits and attention-deficit/hyperactivity disorder (ADHD) on academic outcomes in children. Journal of Consulting and Clinical Psychology, 72(5), 757–766. 10.1037/0022-006X.72.5.757

Biffen, S. C., Warton, C. M. R., Dodge, N. C., Molteno, C. D., Jacobson, J. L., Jacobson, S. W., & Meintjes, E. M. (2020). Validity of automated FreeSurfer segmentation compared to manual tracing in detecting prenatal alcohol exposure-related subcortical and corpus callosal alterations in 9- to 11-year-old children. NeuroImage: Clinical, 28, 102368. 10.1016/j.nicl.2020.102368

Borella, E., Carretti, B., & Pelegrina, S. (2010). The specific role of inhibition in reading comprehension in good and poor comprehenders. Journal of Learning Disabilities, 43(6), 541–552. 10.1177/0022219410371676

Brault, M. C., & Lacourse, É. (2012). Prevalence of prescribed attention-deficit hyperactivity disorder medications and diagnosis among Canadian preschoolers and school-age children: 1994-2007. Canadian Journal of Psychiatry, 57(2), 93–101. 10.1017/CBO9781107415324.004

Bünger, A., Urfer-Maurer, N., & Grob, A. (2021). Multimethod Assessment of Attention, Executive Functions, and Motor Skills in Children With and Without ADHD: Children’s Performance and Parents’ Perceptions. Journal of Attention Disorders, 25(4), 596–606. 10.1177/1087054718824985

Chan, T., Wang, I., & Ybarra, O. (2021). Leading and managing the workplace: The role of executive functions. Academy of Management Perspectives, 35(1), 142–164.

Conners, C. K., Pitkanen, J., & Rzepa, S. R. (2011). Conners 3rd Edition (Conners 3; Conners 2008). In J. S. Kreutzer, J. DeLuca, & B. Caplan (Eds.), Encyclopedia of Clinical Neuropsychology (pp. 675–678). Springer New York. 10.1007/978-0-387-79948-3_1534

Conners, K. (2014). Conners Continuous Performance Test, 3rd Edition. Multi Health Systems.

Conners, Keith, C. (2008). Conners 3rd Edition. Multi Health Systems.

Cortés Pascual, A., Moyano Muñoz, N., & Quílez Robres, A. (2019). The Relationship Between Executive Functions and Academic Performance in Primary Education: Review and Meta-Analysis. Frontiers in Psychology, 10, 1582. 10.3389/fpsyg.2019.01582

Cortese, S., & Coghill, D. (2018). Twenty years of research on attention-deficit/hyperactivity disorder (ADHD): Looking back, looking forward. Evidence-Based Mental Health, 21(4), 173–176. 10.1136/ebmental-2018-300050

Cortese, S., Ph, D., Kelly, C., Chabernaud, C., Martino, A. D., Milham, M. P., & Xavier, F. (2014). Towards systems neuroscience of ADHD: A meta-analysis of 55 fMRI studies. American Journal of Psychiatry, 169(10). 10.1176/appi.ajp.2012.11101521.Towards

Dale, A. M., Fischl, B., & Sereno, M. I. (1999). Cortical surface-based analysis: I. Segmentation and surface reconstruction. Neuroimage, 9(2), 179–194.

Damiani, S., Tarchi, L., Scalabrini, A., Marini, S., Provenzani, U., Rocchetti, M., Oliva, F., & Politi, P. (2021). Beneath the surface: Hyper-connectivity between caudate and salience regions in ADHD fMRI at rest. European Child & Adolescent Psychiatry, 30(4), 619–631. 10.1007/s00787-020-01545-0

Delis, D. C., Kaplan, E., & Kramer, J. H. (2001). The Delis–Kaplan Executive Function System. Pearson Education.

Dewey, D., & Volkovinskaia, A. (2018). Health-related quality of life and peer relationships in adolescents with developmental coordination disorder and attention-deficit-hyperactivity disorder. Developmental Medicine and Child Neurology, 60(7), 711–717. 10.1111/dmcn.13753

Dewey, J., Hana, G., Russell, T., Price, J., McCaffrey, D., Harezlak, J., Sem, E., Anyanwu, J. C., Guttmann, C. R., Navia, B., Cohen, R., Tate, D. F., & Consortium, H. I. V. N. (2010). Reliability and validity of MRI-based automated volumetry software relative to auto-assisted manual measurement of subcortical structures in HIV-infected patients from a multisite study. NeuroImage, 51(4), 1334–1344. 10.1016/j.neuroimage.2010.03.033

Diamond, A. (2013). Executive Functions. Annual Review of Psychology, 64(1), 135–168. 10.1146/annurev-psych-113011-143750

Doebel, S. (2020). Rethinking Executive Function and Its Development. Perspectives on Psychological Science, 15(4), 942–956. 10.1177/1745691620904771

Doom, J. R., Rozenman, M., Fox, K. R., Phu, T., Subar, A. R., Seok, D., & Rivera, K. M. (2021). The transdiagnostic origins of anxiety and depression during the pediatric period: Linking NIMH research domain criteria (RDoC) constructs to ecological systems. Development and Psychopathology, 33(5), 1599–1619. DOI10.1017/S0954579421000559

Edden, R. A. E., Crocetti, D., Zhu, H., Gilbert, D. L., & Mostofsky, S. H. (2012). Reduced GABA concentration in attention-deficit/hyperactivity disorder. Archives of General Psychiatry, 69(7), 750–753. 10.1001/archgenpsychiatry.2011.2280

Espinet, S. D., Graziosi, G., Toplak, M. E., Hesson, J., & Minhas, P. (2022). A Review of Canadian Diagnosed ADHD Prevalence and Incidence Estimates Published in the Past Decade. Brain Sciences, 12(8), 1051. 10.3390/brainsci12081051

Faraone, S. V., Rostain, A. L., Blader, J., Busch, B., Childress, A. C., Connor, D. F., & Newcorn, J. H. (2019). Practitioner Review: Emotional dysregulation in attention-deficit/hyperactivity disorder—Implications for clinical recognition and intervention. Journal of Child Psychology and Psychiatry, and Allied Disciplines, 60(2), 133–150. 10.1111/jcpp.12899

Fischl, B., & Dale, A. M. (2000). Measuring the thickness of the human cerebral cortex from magnetic resonance images. Proceedings of the National Academy of Sciences, 97(20), 11050–11055. 10.1073/pnas.200033797

Fischl, B., Salat, D. H., Busa, E., Albert, M., Dieterich, M., Haselgrove, C., van der Kouwe, A., Killiany, R., Kennedy, D., Klaveness, S., Montillo, A., Makris, N., Rosen, B., & Dale, A. M. (2002). Whole brain segmentation: Automated labeling of neuroanatomical structures in the human brain. Neuron, 33(3), 341–355. 10.1016/s0896-6273(02)00569-x

Fischl, B., van der Kouwe, A., Destrieux, C., Halgren, E., Ségonne, F., Salat, D. H., Busa, E., Seidman, L. J., Goldstein, J., Kennedy, D., Caviness, V., Makris, N., Rosen, B., & Dale, A. M. (2004). Automatically parcellating the human cerebral cortex. Cerebral Cortex (New York, N.Y.L: 1991), 14(1), 11–22. 10.1093/cercor/bhg087

Gau, S. S., Tseng, W.-L., Tseng, W.-Y. I., Wu, Y.-H., & Lo, Y.-C. (2015). Association between microstructural integrity of frontostriatal tracts and school functioning: ADHD symptoms and executive function as mediators. Psychological Medicine, 45(3), 529–543. 10.1017/S0033291714001664

Gioia, G. A., Isquith, P. K., & Guy, Steven. C. (2015). Behavior Rating Inventory of Executive Function®, Second Edition. WPS.

Hai, T., Duffy, H., Lemay, J.-F., Swansburg, R., Climie, E. A., & MacMaster, F. P. (2020). Neurochemical correlates of executive function in children with attention-deficit/hyperactivity disorder. Journal of the Canadian Academy of Child and Adolescent Psychiatry = Journal de l’Academie Canadienne de Psychiatrie de l’enfant et de l’adolescent, 29(1), 15–25.

Hai, T., Swansburg, R. M., Kahl, C., Frank, H., Stone, K. D., Lemay, J.-F., & MacMaster, F. P. (2022). Right Superior Frontal Gyrus Cortical Thickness in Pediatric Attention-Deficit/Hyperactivity Disorder (ADHD). Journal of Attention Disorders, In Press.

Hakkaart-van Roijen, L., Zwirs, B. W. C., Bouwmans, C., Tan, S. S., Schulpen, T. W. J., Vlasveld, L., & Buitelaar, J. K. (2007). Societal costs and quality of life of children suffering from attention deficient hyperactivity disorder (ADHD). European Child & Adolescent Psychiatry, 16(5), 316–326. 10.1007/s00787-007-0603-6

Harmon, S. L., Groves, N. B., Soto, E. F., Sarver, D. E., Irwin, L. N., & Kofler, M. J. (2018). Executive Functioning Heterogeneity in Pediatric ADHD. Journal of Abnormal Child Psychology, 47(2), 273–286. 10.1007/s10802-018-0438-2

Hoogman, M., Bralten, J., Hibar, D. P., Mennes, M., Zwiers, M. P., Schweren, L. S. J., van Hulzen, K. J. E., Medland, S. E., Shumskaya, E., Jahanshad, N., Zeeuw, P. de, Szekely, E., Sudre, G., Wolfers, T., Onnink, A. M. H., Dammers, J. T., Mostert, J. C., Vives-Gilabert, Y., Kohls, G., … Franke, B. (2017). Subcortical brain volume differences in participants with attention deficit hyperactivity disorder in children and adults: A cross-sectional mega-analysis. The Lancet Psychiatry. 10.1016/S2215-0366(17)30049-4

Hoogman, M., Muetzel, R., Guimaraes, J. P., Shumskaya, E., Mennes, M., Zwiers, M. P., Jahanshad, N., Sudre, G., Wolfers, T., Earl, E. A., Soliva Vila, J. C., Vives-Gilabert, Y., Khadka, S., Novotny, S. E., Hartman, C. A., Heslenfeld, D. J., Schweren, L. J. S., Ambrosino, S., Oranje, B., … Franke, B. (2019). Brain Imaging of the Cortex in ADHD: A Coordinated Analysis of Large-Scale Clinical and Population-Based Samples. Am. J. Psychiatry, 176(7), 531–542. 10.1176/appi.ajp.2019.18091033

Huang-Pollock, C. L., Karalunas, S. L., Tam, H., & Moore, A. N. (2012). Evaluating vigilance deficits in ADHD: A meta-analysis of CPT performance. Journal of Abnormal Psychology, 121(2), 360–371. 10.1037/a0027205

Jacobson, L. A., Pritchard, A. E., Koriakin, T. A., Jones, K. E., & Mahone, E. M. (2016). Initial examination of the BRIEF-2 in clinically referred children with and without ADHD symptoms. Journal of Attention Disorders, 1087054716663632. 10.1177/1087054716663632

Joyner, K. B., Silver, C. H., & Stavinoha, P. L. (2009). Relationship Between Parenting Stress and Ratings of Executive Functioning in Children With ADHD. Journal of Psychoeducational Assessment, 27(6), 452–464. 10.1177/0734282909333945

Kahl, C. K., Swansburg, R., Hai, T., Wrightson, J. G., Bell, T., Lemay, J.-F., Kirton, A., & MacMaster, F. P. (2022). Differences in neurometabolites and transcranial magnetic stimulation motor maps in children with attention-deficit/hyperactivity disorder. Journal of Psychiatry and Neuroscience, 47(4), E239–E249. 10.1503/jpn.210186

Kofler, M. J., Irwin, L. N., Soto, E. F., Groves, N. B., Harmon, S. L., & Sarver, D. E. (2019). Executive functioning heterogeneity in pediatric ADHD. Journal of Abnormal Child Psychology, 47(2), 273–286. 10.1007/s10802-018-0438-2

Kofler, M. J., Rapport, M. D., Bolden, J., Sarver, D. E., Raiker, J. S., & Alderson, R. M. (2011). Working memory deficits and social problems in children with ADHD. Journal of Abnormal Child Psychology, 39(6), 805–817. 10.1007/s10802-011-9492-8

MacMaster, F. P., Carrey, N., Sparkes, S., & Kusumakar, V. (2003). Proton spectroscopy in medication-free pediatric attention-deficit/hyperactivity disorder. Biological Psychiatry, 53(2), 184–187. 10.1016/S0006-3223(02)01401-4

Matza, L. S., Paramore, C., & Prasad, M. (2005). A review of the economic burden of ADHD. Cost Effectiveness and Resource AllocationL: C/E, 3, 5. 10.1186/1478-7547-3-5

McAuley, T., & White, D. (2011). A Latent Variables Examination of Processing Speed, Response Inhibition, and Working Memory during Typical Development. Journal of Experimental Child Psychology, 108(3), 453–468. 10.1016/j.jecp.2010.08.009

Miyake, A., & Friedman, N. P. (2012). The Nature and Organization of Individual Differences in Executive Functions: Four General Conclusions. Current Directions in Psychological Science, 21(1), 8–14. 10.1177/0963721411429458

Nigg, J. T., Willcutt, E. G., Doyle, A. E., & Sonuga-Barke, E. J. S. (2005). Causal heterogeneity in attention-deficit/hyperactivity disorder: Do we need neuropsychologically impaired subtypes? Biological Psychiatry, 57(11), 1224–1230. 10.1016/j.biopsych.2004.08.025

Overmeyer, S., Bullmore, E. T., Sucking, J., Smmons, A., Williams, S. C. R., Santosh, P. J., & Taylor, E. (2001). Distributed grey and white matter deficits in hyperkinetic disorder: MRI evidence for anatomical abnormality in an attentional network. Psychological Medicine, 31(8), 1425–1435. 10.1017/s0033291701004706

Polanczyk, G. V., Willcutt, E. G., Salum, G. A., Kieling, C., & Rohde, L. A. (2014). ADHD prevalence estimates across three decades: An updated systematic review and meta-regression analysis. International Journal of Epidemiology, 43(2), 434–442. 10.1093/ije/dyt261

Rapport, M. D., Alderson, R. M., Kofler, M. J., Sarver, D. E., Bolden, J., & Sims, V. (2008). Working memory deficits in boys with attention-deficit/hyperactivity disorder (ADHD): The contribution of central executive and subsystem processes. Journal of Abnormal Child Psychology, 36(6), 825–837. 10.1007/s10802-008-9215-y

Rohr, C. S., Bray, S. L., & Dewey, D. M. (2021). Functional connectivity based brain signatures of behavioral regulation in children with ADHD, DCD, and ADHD-DCD. Development and Psychopathology, 1–10. DOI: 10.1017/S0954579421001449

Rubia, K. (2018). Cognitive neuroscience of attention deficit hyperactivity disorder (ADHD) and its clinical translation. Frontiers in Human Neuroscience, 12(March), 1–23. 10.3389/fnhum.2018.00100

Samea, F., Soluki, S., Nejati, V., Zarei, M., Cortese, S., Eickhoff, S. B., Tahmasian, M., & Eickhoff, C. R. (2019). Brain alterations in children/adolescents with ADHD revisited: A neuroimaging meta-analysis of 96 structural and functional studies. Neuroscience & Biobehavioral Reviews, 100, 1–8. 10.1016/j.neubiorev.2019.02.011

Schneider, H., Ryan, M., & Mahone, E. M. (2020). Parent versus teacher ratings on the BRIEF-preschool version in children with and without ADHD. Child NeuropsychologyL: A Journal on Normal and Abnormal Development in Childhood and Adolescence, 26(1), 113–128. 10.1080/09297049.2019.1617262

Schwörer, M. C., Reinelt, T., Petermann, F., & Petermann, U. (2020). Influence of executive functions on the self-reported health-related quality of life of children with ADHD. Quality of Life Research, 29(5), 1183–1192. 10.1007/s11136-019-02394-4

Shang, C. Y., Wu, Y. H., Gau, S. S., & Tseng, W. Y. (2013). Disturbed microstructural integrity of the frontostriatal fiber pathways and executive dysfunction in children with attention deficit hyperactivity disorder. Psychological Medicine, 43(5), 1093–1107. 10.1017/S0033291712001869

Shaw, P., Eckstrand, K., Sharp, W., Blumenthal, J., Lerch, J. P., Greenstein, D., Clasen, L., Evans, A., Giedd, J., & Rapoport, J. L. (2007). Attention-deficit/hyperactivity disorder is characterized by a delay in cortical maturation. Proceedings of the National Academy of Sciences of the United States of America, 104(49), 19649–19654. 10.1073/pnas.0707741104

Shaw, P., Stringaris, A., Nigg, J., & Leibenluft, E. (2014). Emotion Dysregulation in Attention Deficit Hyperactivity Disorder. American Journal of Psychiatry, 171(3), 276–293. 10.1176/appi.ajp.2013.13070966

Sheehan, D. V., Sheehan, K. H., Shytle, R. D., Janavs, J., Bannon, Y., Rogers, J. E., Milo, K. M., Stock, S. L., & Wilkinson, B. (2010). Reliability and validity of the mini international neuropsychiatric interview for children and adolescents (MINI-KID). The Journal of Clinical Psychiatry, 71(3), 313–326. 10.4088/JCP.09m05305whi

Stern, A., Pollak, Y., Bonne, O., Malik, E., & Maeir, A. (2013). The Relationship Between Executive Functions and Quality of Life in Adults With ADHD. Journal of Attention Disorders, 21(4), 323–330. 10.1177/1087054713504133

Tafazoli, S., O’Neill, J., Bejjani, A., Ly, R., Salamon, N., McCracken, J. T., Alger, J. R., & Levitt, J. G. (2013). 1H MRSI of middle frontal gyrus in pediatric ADHD. Journal of Psychiatric Research, 47(4), 505–512. 10.1016/j.jpsychires.2012.11.011

Toplak, M. E., Bucciarelli, S. M., Jain, U., & Tannock, R. (2009). Executive functions: Performance-based measures and the behavior rating inventory of executive function (BRIEF) in adolescents with attention deficit/hyperactivity disorder (ADHD). Child Neuropsychology, 15(1), 53–72. 10.1080/09297040802070929

Wechsler, D. (2014). WISC-V: Technical and Interpretive Manual. Pearson Education.

Willcutt, E. G., Doyle, A. E., Nigg, J. T., Faraone, S. V., & Pennington, B. F. (2005). Validity of the executive function theory of attention-deficit/hyperactivity disorder: A meta-Analytic review. Biological Psychiatry, 57(11), 1336–1346. 10.1016/j.biopsych.2005.02.006

Wolraich, M. L., Hagan, J. F., Allan, C., Chan, E., Davison, D., Earls, M., Evans, S. W., Flinn, S. K., Froehlich, T., Frost, J., Holbrook, J. R., Lehmann, C. U., Lessin, H. R., Okechukwu, K., Pierce, K. L., Winner, J. D., & Zurhellen, W. (2019). Clinical practice guideline for the diagnosis, evaluation, and treatment of attention-deficit/hyperactivity disorder in children and adolescents. Pediatrics, 144(4). 10.1542/peds.2019-2528

Yang, X., Carrey, N., Bernier, D., & MacMaster, F. P. (2015). Cortical thickness in young treatment-naive children with ADHD. Journal of Attention Disorders, 19(11), 925–930. 10.1177/1087054712455501

